# Investigation of Intestinal Bacteria-binding Monoclonal Antibodies Derived from Rabbit Single B Cells

**DOI:** 10.1101/2024.07.26.605397

**Authors:** Khairil Anwar, Shota Kaneko, Itsuki Kawakita, Daffa Sean Adinegoro, Monami Kihara, Teruyo Ojima-Kato, Hideo Nakano

## Abstract

Immunoglobulins in the intestine have been reported to play a pivotal role in regulating the composition and function of intestinal microbiota. However, the relationship between monoclonal antibodies (mAbs) and intestinal bacteria remains elusive due to the lack of analytical methods. Here we report the utilization of Ecobody technology, a single B cell technique developed by our group, for obtaining and investigating intestinal bacteria-binding mAbs derived from single B cells of a non-immunized rabbit. *Bacteroides cellulosilyticus* isolated from the rabbit stool was used as the antigen for selecting B cells, followed by antibody gene amplification, mAb expression in cell-free protein synthesis (CFPS), and activity evaluation. For large-scale and cost-effective production of the selected mAb, we employed the secretion expression system for antigen-binding fragment (Fab) using *Brevibacillus choshinensis*, and successfully yielded ∼2.0 mg/L-culture of purified antibody with high affinity to the target bacterium. Purified Fab showed binding activity towards various intestinal bacteria, suggesting the broad specificity toward intestinal bacteria-associated antigens.

## 1. Introduction

The gastrointestinal tract (GIT) of both humans and animals is inhabited by a dynamic ecosystem of a million microorganisms, referred to collectively as the intestinal microbiota, which are known to be involved in various activities of the host, including nutrient metabolism, physiological homeostasis and maturation of the mucosal immune system [1, 2]. Intestinal bacteria consist of several species, and their composition differs significantly among healthy individuals [2, 3]. The abnormal alterations of intestinal bacteria may potentially be harmful and result in several diseases, including metabolic [4–6] and immune disorders [7]. Immunoglobulins in the intestine play a pivotal role in facilitating the interaction between host and intestinal bacteria and serve as key effectors that participate in regulating the composition of intestinal bacteria [8–10]. Despite their fundamental role in regulating intestinal microbiota, the specificity of naturally homeostatic antibodies is still elusive and requires further investigation.

A recent technique for monoclonal antibody (mAb) screening and production based on single-cell methods has been established that enables the investigation of the functional properties of intestinal bacteria-binding monoclonal antibodies. The utilization of single-cell techniques allows the comprehensive characterization of antibody diversity in responses to intestinal bacteria and the evaluation of the specificity and affinity of antibodies [8]. Furthermore, studies of the mAbs against intestinal bacteria derived from individual B cells or plasma cells could unravel the gut-immunity mechanism [11], host-microbiota relationships [12, 13] and the development of antibody-based therapeutics for gut-related disorders [14, 15]. More recently, Ecobody technology, developed by our group, is a rapid and cost-effective technique for screening and generating mAbs from single B cells and is suitable for investigating mAbs against intestinal bacteria [16–18].

In this study, we report the investigation of intestinal bacteria-binding monoclonal antibodies derived from single B cells of a non-immunized rabbit by Ecobody technology, a single B cell technology developed by our group as previously reported [17, 18]. *Bacteroides cellulosilyticus* (*B. cellulosilyticus*) isolated from the rabbit stool was used as the antigen for selecting B cells, followed by antibody gene amplification, mAb expression in cell-free protein synthesis (CFPS), and activity evaluation. We adopted the secretion expression system of the antigen-binding fragment (Fab) by *Brevibacillus choshinensis* for large-scale and low-cost antibody production. *Brevibacillus choshinensis*, belonging to soil-derived Gram-positive bacteria, was first isolated by Takagi *et al.* [19] and discovered to naturally secrete significant quantities of proteins with low protease and endotoxin activity [20]. In addition, this system possesses several merits because of its robust secretion machinery and lower costs compared to mammalian expression systems.

Using this system, we successfully produced mAb with high affinity to the target bacterium. Our findings demonstrate that the purified Fab cross-reacted toward several intestinal bacteria but did not exhibit binding activity to unrelated antigens, suggesting the specificity toward intestinal bacteria-associated antigens. This binding included various Gram-negative and Gram-positive bacteria. Interestingly, the Fab also targeted lipopolysaccharide (LPS) extracted from Gram-negative bacteria. Therefore, these findings may provide valuable insights for understanding the relationship between the host immune system and the intestinal microbiota.

## 2. Materials and Methods

### 2.1. Isolation of intestinal bacteria from a non-immunized rabbit

Faecal samples were collected from the cecum of a non-immunized rabbit (New Zealand White rabbit, 13 weeks old) and 8.4 mg of the stool was suspended in 1 mL of phosphate-buffered saline (PBS). The diluted faecal solution was seeded on GAM agar medium (Nissui, Japan). The plates were anaerobically incubated at 37°C for 48 hours. The 16S rDNA gene from the obtained colonies was amplified by colony PCR using forward universal primer 7F (5’-AGAGTTTGATYMTGGCTCAG-3’) and reverse universal primer 1510R (5’-ACGGYTACCTTGTTACGACTT-3’). 20 µL of a total reaction mixture contained 1.0 µL of the diluted colony as the template, 2.0 µL of 10× Ex Taq buffer (20 mM of Mg^2+^ plus), 0.2 mM of dNTPs, 1.25 U of Ex Taq DNA polymerase (Takara, Japan) and 0.5 µM of each primer. The PCR was run using the following program: 2 min at 94°C; 25 cycles of 15 s at 98°C, 30 s at 50°C, and 1 minute 40 s at 72°C; and 1 min at 72°C. A single band was confirmed at the position of approximately 1.5 kbp, which is almost the entire length of 16S rDNA genes. The sequencing analysis of 16S rDNA genes was subsequently performed to reveal bacterial species. The 16S rDNA sequences were compared with a genetic database to identify bacterial species. *Bacteroides cellulosilyticus* and *Parabacteroides distasonis* were identified from the rabbit faecal sample. Therefore, *B. cellulosilyticus* was used as the antigen in this study. This study was approved by the Committee of Animal Experiments of the Graduate School of Bioagricultural Sciences, Nagoya University (permit number: 2018053101) and carried out according to the Regulations on Animal Experiments at Nagoya University.

### 2.2. Analysis of rabbit intestinal microbiota

Bacterial genomic DNA was extracted from the collected faecal sample using the NucleoSpin DNA stool (Takara, Japan). PCR was performed to prepare 16S rDNA libraries using primers with adapter sequences for analysis. The primers used were 16S-Ilumina-F (5’-TCGTCGGCAGCGTCAGATGTGTATAAGAGACAGCCTACGGG NGGCWGCAG-3’) and 16S-Ilumina-R (5’-GTCTCGTGGGCTCGGAGATGTGTA TAAGAGACAGGACTACHVGGGTATCTAATCC-3’). PCR was performed in a total reaction volume of 50 µL containing 1.0 µL of bacterial genomic DNA (44.3 ng of gDNA), 10 µL of 5× PrimeSTAR Buffer (Mg^2+^ plus), 0.2 mM of dNTPs, 0.5 U of PrimeSTAR HS DNA Polymerase (Takara, Japan), 0.2 µM of each primer. The PCR reaction program used was as follows: 3 min at 95°C; 25 cycles of 30 s at 95°C, 30 s at 55°C, and 60 s at 72°C; and 3 min at 72°C. Sequence analysis of the prepared DNA library was carried out using MiSeq (Illumina, Inc.) by Gifu University NGS Service.

### 2.3. Preparation of antigens

The isolated *B. cellulosilyticus* was inoculated into 10 mL of liquid GAM medium (Nissui, Japan) and anaerobically incubated at 37°C for 48 hours. The cell pellet was collected by centrifugation, rinsed with PBS, treated with 0.5% formalin in PBS at 80°C, and then stored at -20°C. Lipopolysaccharide (LPS) was also extracted from *B. cellulosilyticus* using LPS Extraction Kit (Intron Biotechnology, South Korea) following the protocol as recommended by the manufacturer.

### 2.4. Isolation of B cells

A few mL of blood samples were collected from the peripheral blood of a non-immunized rabbit. A fraction of lymphocytes was separated using density-gradient centrifugation as previously described in the reference [16]. The lymphocytes including B cells in the middle layer were obtained and rinsed with PBS twice by two rounds of centrifugation (500 × *g*, 20 min, 18°C). After discarding the supernatant, precipitated cells were resuspended in 2 mL of Bun Bunker (Nippon Genetics, Japan), and 1 mL was dispensed to serve as the lymph cell preservation solution. The number of cells was determined using a microscope (Olympus M021, Olympus) with a Bürker-Turk hemocytometer (Japan Clinical Instruments, Japan) and immediately used for further experiment in Section 2.5.

### 2.5. Screening of antigen-specific B cells

The inactivated *B. cellulosilyticus* cells with 0.5% formalin isolated from the rabbit cecum were used for screening antigen-specific B cells. Inactivated cells of *B. cellulosilyticus* (OD_600 nm_ = 0.1) and 1% bovine serum albumin (BSA) were coated on the MaxiSorp plate (Thermo Fisher Scientific), followed by overnight incubation at 4°C. Subsequently, 100 µL of the lymphoid cell fraction as described above was applied to each well in which 1% BSA was immobilized, and incubated at room temperature for 1 hour. The supernatant was collected by gently pipetting to the side of the well after non-specific cells were bound to BSA. The recovered supernatant was subsequently added to each well in which the inactivated cells of *B. cellulosilyticus* were immobilized, and incubated at room temperature for 1 hour. After removing the supernatant and washing twice with PBS, papain solution (PBS pH 7.4 containing 0.25% papain, 0.5% cysteine) was added to obtain B cells bound to *B. cellulosilyticus* cells. The B cell-contained supernatant was collected after incubation for 1 hour. After centrifuging and resuspending in 100 µL of PBS, the f.sight (Cytena) was used to adjust the size to 3 µm to 14 µm and circularity to 0.4 to 1.0. Lymphocytes were collected one by one and divided into a 384-well PCR/QPCR plate (Nippon Genetics, Japan) containing 5 µL of PBS.

### 2.6. Antibody gene amplification from rabbit single B cells

Antibody genes were amplified from rabbit single B cells using specific primers following the primer sequences as described previously in the reference [17]. Additionally, specific primers for the amplification of rabbit IgA were shown in Supplementary Table 1. Single-cell RT-PCR was conducted using SuperScript IV reverse transcriptase (Life Technologies) in a 10 µL reaction mixture containing 0.2 µM of each specific primer. The RT reaction was set according to the protocol recommended by the manufacturer and then added to each well containing a single B cell, followed by incubation at 50°C for 15 minutes. Subsequently, the RT-PCR products were directly utilized for two-step PCR to amplify the heavy chain (Hc) and light chain (Lc) genes independently. The 1^st^ PCR was carried out with a final volume of 10 µL containing 1.0 µL of RT-PCR product, 5.0 µL of 2× buffer for KOD FX Neo, 0.2 mM of dNTPs, 0.2 U of KOD FX Neo polymerase (all purchased from Toyobo, Japan), and mix of primers (0.5 µM of each primer for 1^st^ PCR). The 1^st^ PCR was run using the following protocol: 2 min at 94°C; 30 cycles of 10 s at 94°C, 15 s at 60°C, and 20 s at 68°C; and 3 min at 68°C. Next, the 2^nd^ PCR was performed in a 10 µL reaction volume containing 1.0 µL of 1^st^ PCR products as the templates, 0.25 U of Tks Gflex DNA polymerase (Takara, Japan), reaction buffer (containing Mg^2+^ and dNTP plus) provided by the manufacturer, and mix of primers (0.5 µM of each primer for 2^nd^ PCR). The program for the 2^nd^ PCR was set as follows: 2 min at 94°C; 30 cycles of 10 s at 98°C, 15 s at 60°C, and 20 s at 68°C; and 3 min at 68°C. The 2^nd^ PCR was performed to prepare the connect tails needed for the subsequent Gibson assembly step.

### 2.7. Preparation of DNA fragments for CFPS

The second PCR products were combined by Gibson assembly to pRSET-based cloning vectors which contain a T7 promoter, SKIK (Ser-Lys-Ile-Lys) peptide sequence at the N-terminus, constant region of Hc and Lc, leucine zipper (LZ), PA or HA tag, and T7 terminator. Leucine zippers, LZA and LZB, were fused at the C-terminus of the Hc and Lc, respectively. Linearized pRSET vectors were prepared by PCR with overlapping primers as described in the reference [17], followed by *Dpn*I digestion (Takara, Japan) at 37°C for 1 hour and purification using FastGene Gel/PCR Extraction Kit (Nippon Genetics, Japan). After mixing 1 µL of the linearized vector (100 ng), 1 µL of the second PCR product, and 2 µL of 2× Gibson Master Mix (New England Biolabs, Ipswich, MA), the mixture was incubated at 50°C for 15 min. DNA fragments which include the T7 promoter, antibody gene (Lc or Hc), and T7 terminator for the CFPS were subsequently amplified from the assembled product by PCR with PrimeSTAR Max DNA Polymerase (Takara, Japan). They were directly utilized as the template for the following CFPS.

### 2.8. Cell-free protein synthesis

The modified Fab, also referred to as a zipbody format [17, 18], was synthesized by PURE*frex* system (Gene Frontier, Japan). Oxidized glutathione (GSSG), reduced glutathione (GSH), DsbC, DnaK, and cysteine were added into the reaction solution, as recommended by the manufacturer. The Fab was expressed in a final volume of 10 µL containing 0.5 µL of PCR products or 20 ng of purified DNA fragments (each of Lc and Hc) as the templates, and incubated at 37°C for 90 min using a thermal cycler.

### 2.9. SDS-PAGE for CFPS product analysis

The Fab produced by CFPS was analyzed using SDS-PAGE under reduced conditions following the protocol as previously described in a reference [21]. Briefly, proteins were separated on SDS-PAGE and visualized by Western blotting using horse-radish peroxidase (HRP)-conjugated antibodies (Anti-PA tag Rat Monoclonal Antibody [Fujifilm Wako Pure Chemical Corporation, Japan] for detecting Lc and Anti-HA tag antibody [Novus Biologicals, USA] for detecting Hc) and 1-Step™ Ultra TMB-Blotting Solution (Thermo, Waltham, MA).

### 2.10. Enzyme-linked immunosorbent assay (ELISA)

The binding activity of the synthesized Fab to the antigens was examined by ELISA. Inactivated cells of *B. cellulosilyticus* and 1% BSA were used as the antigens. 50 µL of inactivated cells of bacteria (OD_600 nm_ = 0.1) and 1% BSA diluted in 0.1 M carbonate-bicarbonate buffer, pH 9.6 (Fujifilm Wako Pure Chemical Corporation, Japan) was coated to each well of the MaxiSorp plate (Thermo Fisher Scientific), and incubated overnight at 4 °C. Then, each well was filled with PBS containing 3% skim milk, followed by incubation at 37°C for 1 hour. After washing each well with PBS containing 0.05% tween 20 (PBST) twice, the CFPS product (1^st^ Ab) diluted five times by Can Get Signal Immunoreaction Enhancer Solution 1 (Toyobo, Japan) was added into each well, incubated at 37°C for 1 hour, and washed thrice with PBST. Subsequently, 100 µL of Anti-PA tag Rat Monoclonal Antibody (HRP-conjugated) diluted 5000-fold with Can Get Signal Immunoreaction Enhancer Solution 2 (Toyobo, Japan) was added to each well, incubated at 37°C for 30 min and washed thrice with PBST. 50 µL of ELISA POD Substrate TMB Solution (Nacalai Tesque, Japan) was added to each well and incubated at room temperature for 10 min. After adding 50 µL of 2 M H_2_SO_4_ to stop the reaction, the absorbance at 450 nm was measured with a plate reader (M-200 Pro, Tecan).

### 2.11. Plasmid preparation and DNA sequence analysis

The assembled constructs containing the amplified mAb genes as described in the section 2.7. were transformed into *E. coli* DH5α competent cells (Takara, Japan) On LB-medium plates with 100 µg/mL ampicillin, several colonies were selected for colony PCR and sequencing analysis using the same primer sets as for preparation of the CFPS templates.

### 2.12. Expression of Fab in the *Brevibacillus choshinensis* expression system

The plasmid for Fab in *Brevibacillus* expression was constructed using the Hi-Fi assembly method (New England Biolabs). The original Fab format (without the SKIK tag and LZ tag) was expressed as the secretory protein in *Brevibacillus choshinensis*. The Lc and Hc derived from clone No. 86, which was amplified with the primer sets connecting to both ends, were inserted into linearized pNCMO2 vectors (Takara, Japan), which were amplified with primer sets shown in Supplementary Table 2. The secretion signal 3 from pBIC3 (Takara, Japan) and secretion signal 4 from pBIC4 (Takara, Japan) were incorporated upstream of the Lc and Hc genes, respectively, to allow protein secretion. A polyhistidine tag (6×his tag) was fused downstream of the Lc and Hc genes, respectively. The amplified vectors and insert genes were digested using *Dpn*I enzyme (New England Biolabs), followed by DNA purification and Hi-Fi assembly (New England Biolabs). Subsequently, the Hi-Fi assembled constructs were transformed into *E. coli* DH5α competent cells (Takara, Japan) for plasmid preparation. The plasmid containing correct sequences of the Lc and Hc was extracted from *E. coli* and subsequently transformed into *Brevibacillus choshinensis* HPD31-SP3 competent cells (Takara, Japan) for protein expression following the protocol as recommended by the manufacturer. The plasmid map of pNCMO2-Fab is shown in Supplementary Fig. S1.

TM medium supplemented with 200 mM Arginine Hydrochloride (ArgHCl), 10 g/L of proline, and 60 mM MgSO_4_ was used to culture the transformed *Brevibacillus choshinensis*. The TM medium contained 10 g/L of phytone peptone (Thermo Fisher Scientific), 10 g/L of glucose, 5.75 g/L of 35% Ehrlich bonito extract (Kyokuto Pharmaceutical, Japan), 2 g/L of bacto yeast extract (Thermo Fisher Scientific), 10 mg/L of FeSO_4_·7H_2_O, 8.9 mg/L MnCl_2_·4H_2_O, and 1 mg/L of ZnSO_4_·7H_2_O. Preculture was prepared by inoculating a single colony of the transformed *Brevibacillus choshinensis* into 3 mL of the TM medium containing 50 μg/mL of neomycin and incubated at 37°C for 24 hours with shaking at 110 rpm. Subsequently, 1 mL of preculture was mixed with the supplemented TM medium. The mixture was dispensed into each well of 96-deep-well plates. The plates were covered by a gas permeable seal (Biomedical Science, Japan) and incubated at 30°C for 55 hours with reciprocal shaking (1000 rpm) in a plate incubator (MBR-022UP, TAITEC). After centrifuging at 5,000 g for 15 min, the culture supernatant containing Fab from each well was harvested and sanitized by filtering (0.8 µm filter). Furthermore, the culture supernatant was analyzed by SDS-PAGE under reduced and non-reduced conditions and visualized using Western blotting with Anti-His tag mAb-HRP conjugate. Dithiothreitol (DTT) was used as a reducing agent.

### 2.13. Purification of Fab

The culture supernatant (222 mL) of *Brevibacillus choshinensis* collected from 96-deep-well plates was dialyzed overnight against buffer composed of 100 mM Tris-HCl and 150 mM NaCl (pH 8.0) and diluted twice with binding buffer containing 100 mM Tris-HCl, 500 mM NaCl and 10 mM imidazole (pH 8.0). The diluted supernatant was mixed with 2 mL of Ni-NTA agarose HP (Fujifilm Wako Pure Chemical Corporation, Japan), shaken at 100 rpm (4°C) for 1 hour, and then loaded onto a column. After extensive washing of the column with wash buffer composed of 100 mM Tris-HCl, 500 mM NaCl, and 20 mM imidazole (pH 8.0), the Fab was eluted using elution buffer containing 100 mM Tris-HCl, 500 mM NaCl, and 300 mM imidazole (pH 8.0). Fractions containing the Fab in elution buffer were pooled and dialyzed overnight against the buffer composed of 100 mM Tris-HCl and 200 mM NaCl (pH 8.0). The dialyzed samples were further purified using size exclusion chromatography (SEC) with HiPrep^TM^ 16/60 Sephacryl S-200 HR (Cytiva) that was equilibrated with the buffer containing 30 mM Tris-HCl and 200 mM NaCl (pH 8.0). The monomer fractions were then eluted and used for further experiments. The purified Fab was analyzed by non-reduced SDS-PAGE conditions and visualized by Coomassie Brilliant Blue (CBB) staining.

### 2.14. Cross-reactivity evaluation of Fab

The cross-reactivity of the purified Fab expressed in *Brevibacillus choshinensis* was evaluated by ELISA. Three strains of Gram-negative bacteria (*Parabacteriodes sp.* [RD014198], *Christensenella sp.* [RD014214], *Parasutterella sp.* [RD014256] and their LPS), seven strains of Gram-positive bacteria (*Catabacter sp.* [RD014212], *Clostridium sp.* [RD014215], *Flavonifractor sp.* [RD014238], *Intestinimonas sp.* [RD014241], *Peptoniphilus sp.* [RD014243], *Faecalicoccus sp.* [RD014249], and *Mediterraneibacter sp.* [RD014895]), *S. cerevisiae* and HEK293T cells prepared as in Section 2.3. were used as the antigens. All RD bacterial strains were purchased from the National Institute of Technology and Evaluation (NITE), Kisarazu, Japan. *B. cellulosilyticus* was used as the positive control. In ELISA, 50 µL of inactivated cells (OD_600 nm_ = 0.1) or 10 µg/mL of LPS or 1% BSA in PBS was coated on the MaxiSorp plate (Thermo Fisher Scientific) overnight at 4°C. ELISA was performed following the protocol as previously described in Section 2.10 using the purified Fab (10 µg/mL) produced in *Brevibacillus choshinensis* as the primary antibody and Anti-His tag mAb-HRP conjugate (MBL, Japan) as the secondary antibody.

### 2.15. Kinetics analysis of Fab

The kinetic analysis was conducted based on the biolayer interferometry using BLItz (Pall ForteBio) at room temperature with aminopropyl-silane (APS) sensors as previously described in the reference [17]. The sensors were washed with 400 µL of PBS for 30 s. The inactivated *B. cellulosilyticus* cells (OD_600 nm_ = 1.0) were immobilized on the sensor for 120 s and the sensor was washed with 400 µL of PBS for 120 s. The purified Fab was diluted with 1× kinetics buffer (Pall ForteBio) at concentrations of 20, 50, 100, and 200 mM. 4 µL of the diluted Fab was then associated with the sensor for 300 s, followed by dissociation using 400 µL of PBS for 300 s. The *k*_a_, *k*_d_, and *K*_D_ values were calculated using a global fitting mode in a 1:1 binding model.

## 3. Results

### 3.1. Isolation of intestinal bacteria

We derived the rabbit stool from the cecum in a sterile manner to determine a particular intestinal microbiota that binds to the antibodies. Several colonies were randomly chosen after anaerobic incubation on GAM agar medium. Two isolates were successfully identified as *Bacteroides cellulosilyticus* and *Parabacteroides distasonis* by comparing a partial nucleotide sequence of 16S rDNA genes to the GeneBank database (BLAST NCBI) with 99% sequence similarity for both strains, because organisms are generally regarded as closely related and most likely strains belonging to the same species when they share a sequence identity of ≥97% [22]. As no colony of *Parabacteroides distasonis* grew when inoculated to new plates, we further used *Bacteroides cellulosilyticus* as the antigen for the selection of antigen-specific B cells.

### 3.2. Composition of rabbit intestinal microbiota

The sequencing analysis of 16S rDNA genes was performed to confirm that isolated bacteria are present in the rabbit cecum. The extracted DNA genomic was utilized to prepare a 16S rDNA library for sequencing analysis. The composition of intestinal microbiota is depicted in Fig. 1. Firmicutes were the most abundant phylum, accounting for around 81% of the gut bacteria in the cecum. The second most abundant phylum was Bacteroidetes (>5%), which is the entire class of Bacteroidia, followed by Akkermansia (phylum Verrucomicrobia), accounting for 5% of the total bacteria in the cecum. The phylum Actinobacteria was also detected in a percentage >3%. Euryarchaeota was the only phylum classified under the kingdom Archae that could be identified, but it was present at a relatively very low abundance (0.7%). At the family level, Ruminococcaceae and Lachnospiraceae were found to be overrepresented in a percentage reaching around 30% and 25%, respectively. Additional families with sizable abundance included Eubacteriaceae (15%) and Christensenellaceae (7%). These four families belong to the phylum Firmicutes. The other families, such as Rikenellaceae, Atopobiaceae, Eggerthellaceae, Tennerellaceae, and Bacteroidaceae were classified, but these bacteria were present at relatively low abundance (0.5-3%).

**Fig 1.**
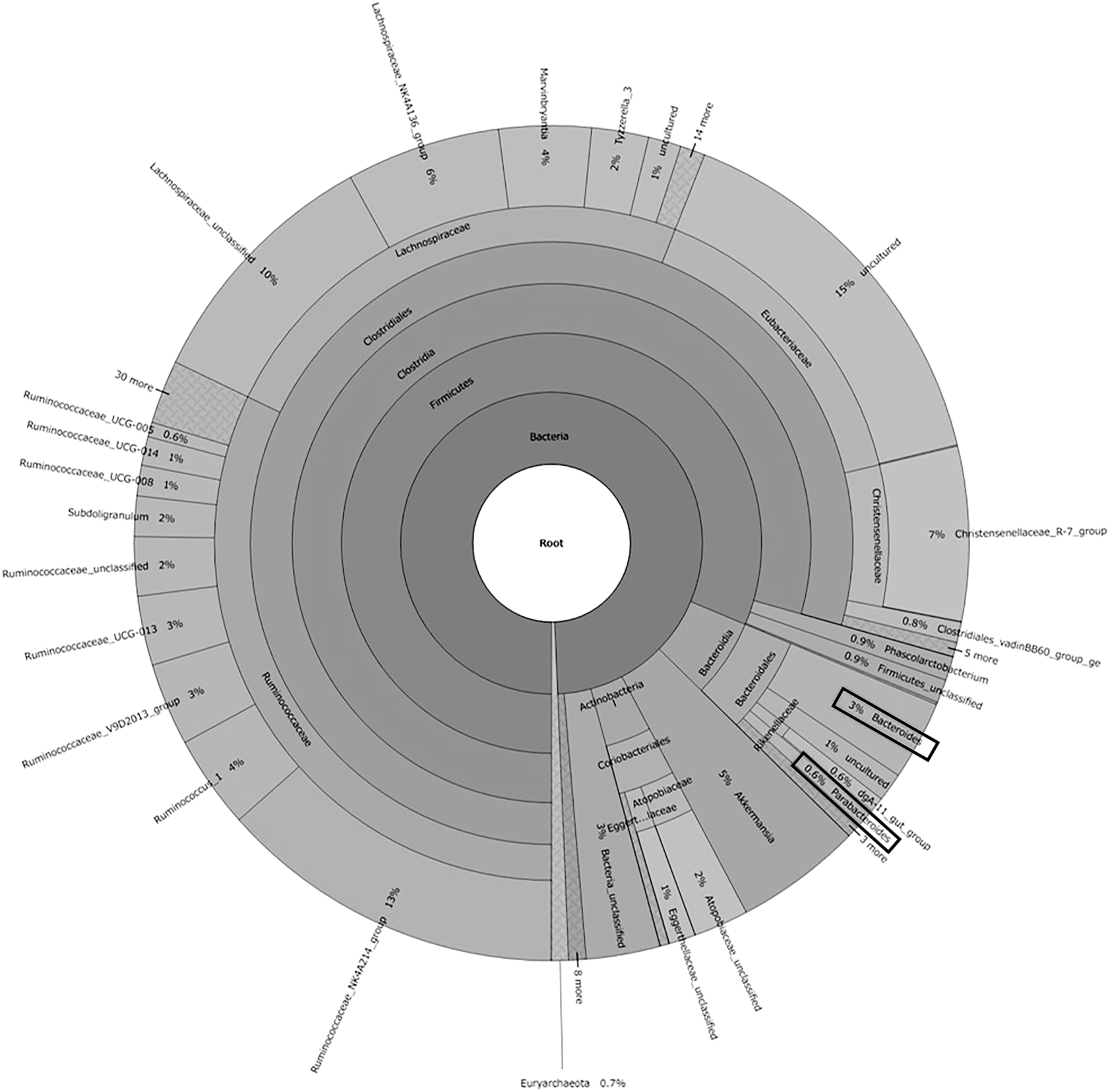
Composition of the rabbit intestinal microbiota based on the result of 16S rDNA gene sequencing. The bacterial genomic DNA was extracted from the fecal sample of the rabbit cecum. The genus *Bacteroides* and *Parabacteroides* obtained in this study are highlighted within a black frame.

The five most abundant genera were the uncultured Eubacteriaceae group (15%), Ruminococcaceae NK4A214 group (13%), unclassified Lachnospiraceae group (10%), Christensenellaceae R-7 group (7%), Lachnospiraceae NK4A136 group (6%). Several genera, including Ruminococcus 1, Marvinbryantia, Ruminococcaceae V9D2013 group, Ruminococcaceae UCG-013, Bacteroides, unclassified Ruminococcaceae group, Subdoligranulum, unclassified Atopobiaceae group and Tyzzerella 3, were found in a relatively low percentage (2-4%). While the smallest relative abundances at the genus level were unclassified Eggerthellaceae group, Ruminococcaceae UCG-008, Ruminococcaceae UCG-014, uncultured Lachnospiraceae group, Phascolarctobacterium, Clostridiales vadin BB60 group, Ruminococcaceae UCG-005, dgA-11 gut group and Parabacteriodes; their proportion ranges from 0.6-1%.

### 3.3. Rabbit mAb screening

The screening of mAb was carried out using Ecobody technology as previously described in reference [17, 18] to generate mAbs that bind to intestinal bacteria. First, 7.5 × 10^4^ lymphocytes were collected from a non-immunized rabbit. Cells that nonspecifically bound to BSA were removed and 1.75 × 10^4^ cells were obtained.

Subsequently, approximately 7.5 × 10^3^ cells bound to the inactivated *B. cellulosilyticus* cells and were treated with Papain solution. 2.3 × 10^3^ cells were obtained after final resuspension in 100 µL of PBS. Single cells were sorted using f.sight (Cytena) and transferred into a PCR well plate, of which 17 cells were used for single-cell PCR to acquire the antibody genes. From them, eight Lc and four Hc genes around 421 bp were successfully obtained after 2^nd^ PCR (Fig. 2). However, only clone No. 86 contained the Lc and Hc genes derived from the same cell. The antibody genes from each clone were named L1, L2, L3, L7, L8, L13, L14, and L86 for Lc genes and M1, M4, M12 and M86 for Hc genes. The IgM type Hc genes were obtained, and no IgG or IgA type Hc was recovered.

**Fig 2.**
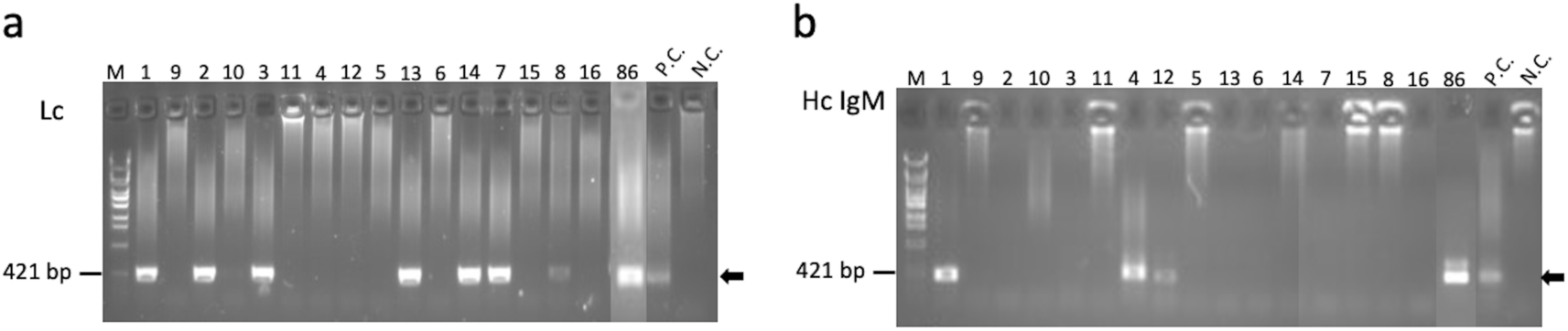
The amplification results of antibody genes after second PCR. (a) Lc genes and (b) Hc IgM genes. P. C. indicates positive control using peripheral blood lymphocyte RNA from which B cells were obtained. N. C. means negative control without B cells. M is DNA size marker λ-EcoT14 I digest. Black arrows indicate the target genes.

Nine Fab clones by combining Lc (L13, L14, and L86) and Hc (M1, M4, and M86) of acquired antibody genes were synthesized in the CFPS and subsequently evaluated by ELISA. The sequences of variable region amino acids of these expressed clones are displayed in Fig. 3. All the clones were expressed as zipbodies with the SKIK tag fused at the N-terminus of the Hc and Lc. Protein expression was confirmed by Western blotting using Anti-PA tag Rat Monoclonal Antibody for detecting Lc and Anti-HA tag antibody for detecting Hc (Supplementary Fig. S2). L13M86 demonstrated the highest binding activity against *B. cellulosilyticus* as the antigen, followed by L13M1 and L86M86 (Fig. 4). Additionally, L86M86, consisting of the cognate pairing of Lc and Hc, showed a reasonably good signal with low cross-reactivity. Consequently, Fab clone L86M86 was selected and produced in the *Brevibacillus choshinensis* expression system for further analyses.

**Fig. 3.**
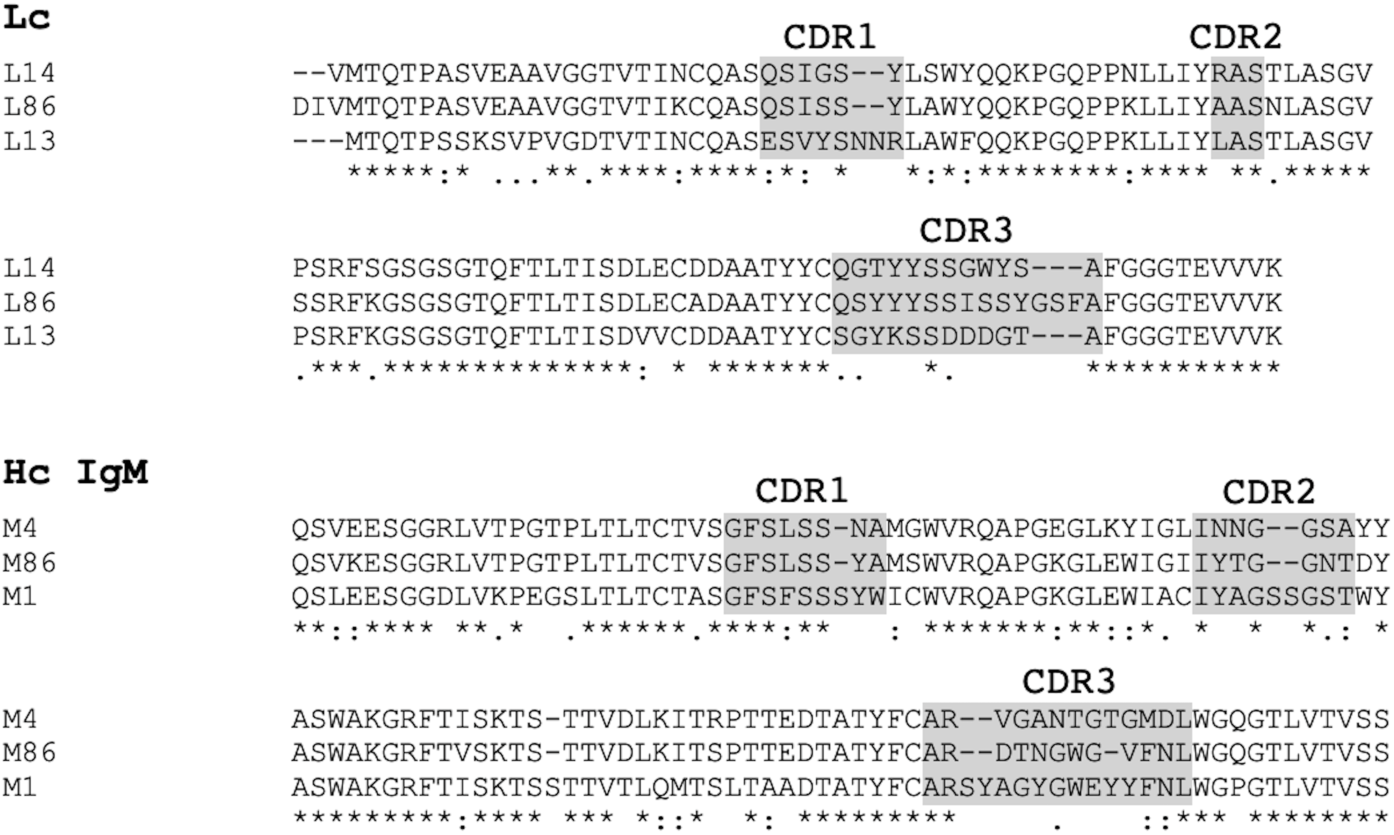
The sequences of variable region amino acids of the expressed antibody clones. Complementarity-determining regions (CDRs) are highlighted in gray.

**Fig. 4.**
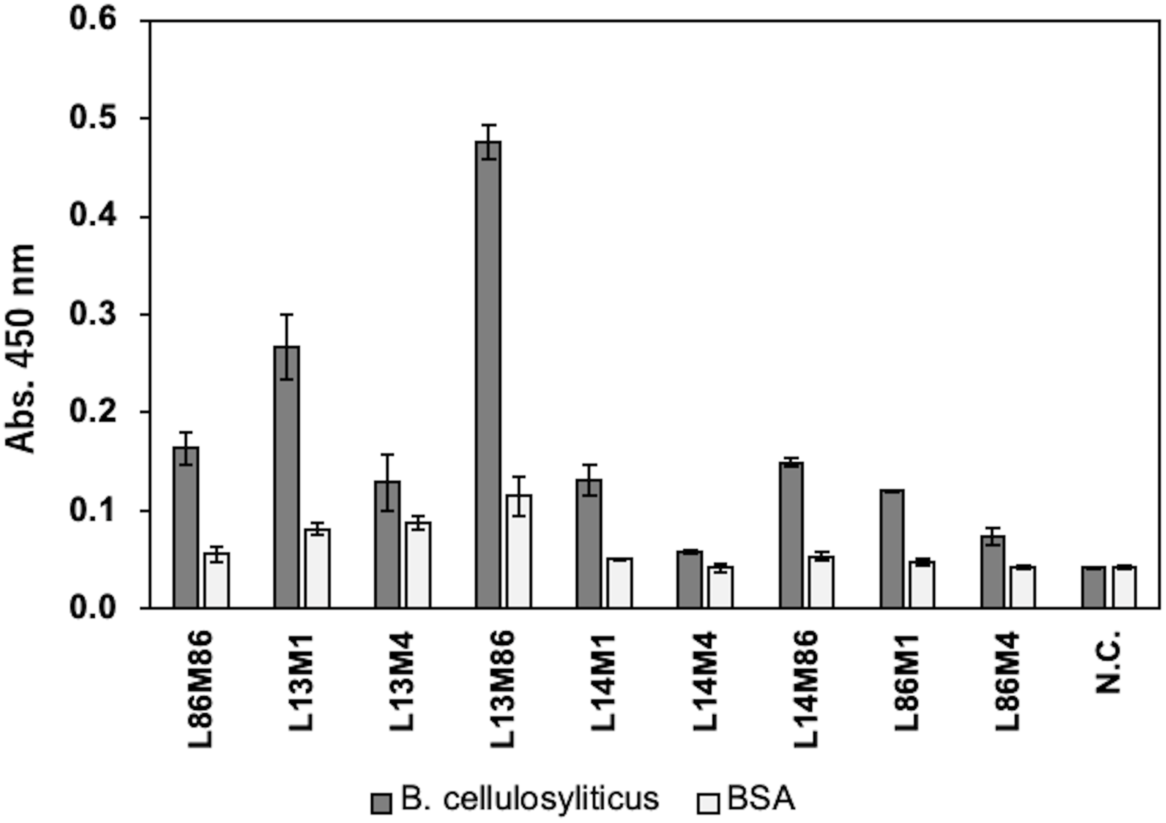
ELISA results against *B. cellulosilyticus* of the synthesized Fab obtained by Ecobody technology. Nine Fab clones by combining Lc (L13, L14, and L86) and Hc (M1, M4, and M86) were synthesized by CFPS. N. C. means the negative control using the CFPS product without DNA templates as the primary antibody. Bars indicate the average of two measurements ± standard deviations.

### 3.4. In vivo expression of selected Fab in the *Brevibacillus choshinensis* expression system

The Fab was successfully produced in *Brevibacillus choshinensis* as the secretory protein to obtain an active form. We found that the TM medium supplemented with 200 mM arginine hydrochloride (ArgHCl), 10 g/L of proline, and 60 mM MgSO_4_ achieved the best condition for antibody expression in *Brevibacillus choshinensis* (data not shown). We decided to use these molecules due to their well-defined role as folding auxiliary molecules and efficiency for various recombinant proteins expressed in *Brevibacillus choshinensis* [23–27]. After determining the medium composition in 3 mL test tube culture, the transformed *Brevibacillus choshinensis* was cultivated at 96 mL scale for antibody production using the plate containing the supplemented TM medium in all wells. The harvested culture supernatant containing Fab was analyzed by SDS-PAGE and visualized by Western blotting with anti-His tag-HRP conjugated antibodies.

The results showed that a single band (around 25 kDa) under reduced conditions corresponding to Hc and Lc proteins was confirmed and presumed to be overlapping Hc and Lc proteins. The Fab protein was also present as a single band (around 45 kDa) under non-reduced conditions, indicating the association between Hc and Lc proteins (Fig. 5a). The culture supernatant and BSA (3.125 – 100 µg/mL) were analyzed by SDS-PAGE to estimate the protein amount. Around 9.4 mg of Fab protein produced per Liter of the culture in 96-deep-well plates was estimated by protein band intensity with CBB staining and Image J quantitation. We used 222 mL of the culture supernatant harvested from the 96-deep-well plates culture containing around 2.71 mg of the Fab and purified by Ni-affinity chromatography, followed by SEC. The SEC chromatogram exhibited a peak around 64-76 mL, suggesting the intact Fab is successfully formed and eluted as a monomer fraction (Fig. 5b). The final yield of purified Fab was approximately 0.57 mg per 222 mL of the culture supernatant as shown in Supplementary Table 3. The protein band pattern of the purified Fab under non-reduced SDS-PAGE conditions was almost the same as before purification, in which clear bands at the expected size were confirmed, as shown in Fig. 5c. These results demonstrated that Fab proteins were successfully purified as the correct association of Hc-Lc.

**Fig. 5.**
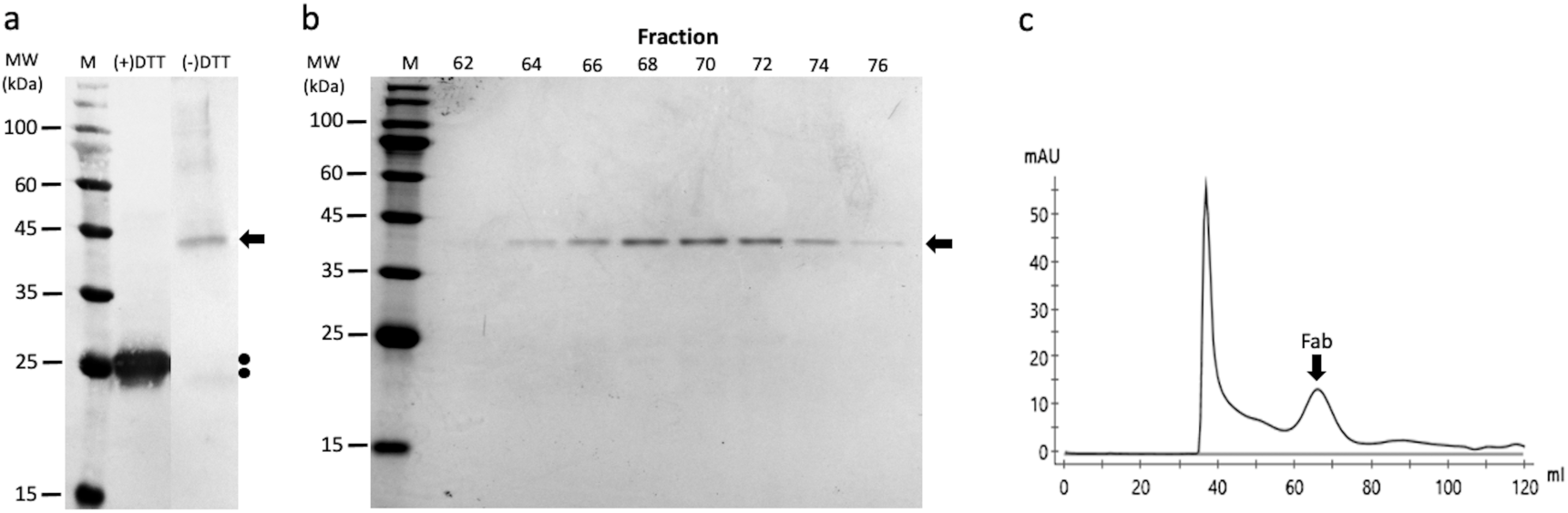
Analysis of Fab produced in the *Brevibacillus choshinensis* expression system. a) Analysis of the culture supernatant using SDS-PAGE under reduced (+DTT) and non-reduced (-DTT) conditions and visualization using Western blotting with Anti-His tag mAb-HRP conjugate. The Fab protein band is shown by the arrow and Hc and Lc proteins are shown by dots. M indicates protein markers. b) CBB staining of the purified Fab after non-reduced SDS-PAGE conditions. The arrow indicates the expected size of the purified Fab. M means protein markers. c) SEC elution profile of the purified Fab. Elution positions of the Fab are shown by the arrow.

### 3.5. Cross-reactivity of the purified Fab against intestinal bacteria

Antigen binding activity of the purified Fab produced in the *Brevibacillus choshinensis* expression system against multiple bacteria was examined by ELISA. In this study, we used commonly distributed human enterobacterial strains to determine the cross-reactivity, since rabbit intestinal bacteria were not widely available. The results showed that Fab had cross-reactivity toward all examined bacteria, however it did not cross-react with BSA. As shown in Fig. 6a, the binding activity of Fab toward inactivated Gram-negative bacterial cells was slightly greater than that of their LPS. Furthermore, we verified that Fab had the ability to attach to Gram-positive bacteria. Fab exhibited a higher ELISA signal when exposed to *Mediterraneibacter sp.* (RD014895) compared to other antigens, including *B. cellulosilyticus*. Contrarily, it did not cross-react with *S. cerevisiae* and HEK293T cells [Fig. 6b], indicating that the Fab did not recognize unrelated antigens with intestinal bacteria. These findings suggested that monoclonal IgM antibodies targeting *B. cellulosilyticus* were functionally active, even though cross-species reactivity against multiple intestinal bacteria was observed, potentially attributed to IgM originating from single B cells of a non-immunized rabbit.

**Fig. 6.**
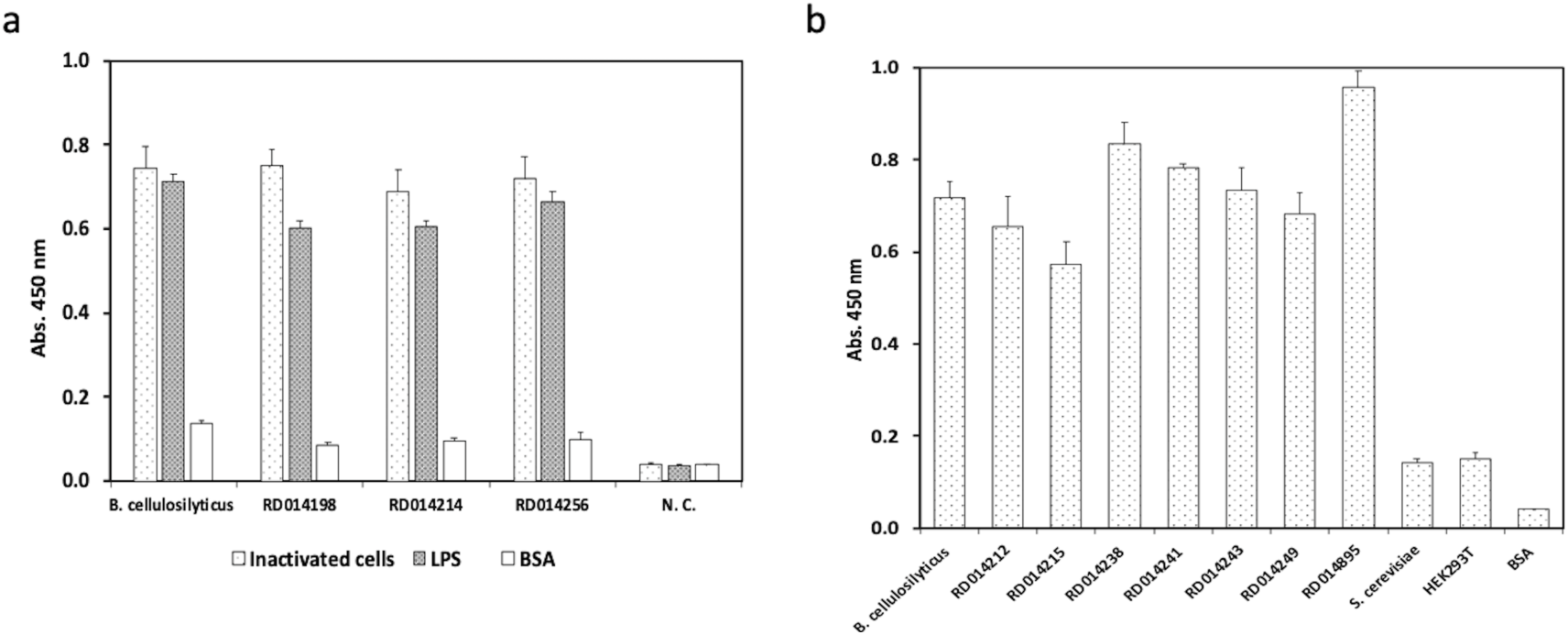
ELISA results of the purified Fab produced in the *Brevibacillus choshinensis* expression system against multiple bacteria. a) The binding activiy of the Fab against Gram-negative bacteria and their LPS. Antigens are *B. cellulosilyticus* (positive control), RD014198 (*Parabacteroides sp.*), RD014214 (*Christensenella sp.*), RD014256 (*Parasutterella sp.*), and BSA. N. C. means negative control using PBS as the primary antibody. b) The binding activiy of the Fab against Gram-positive bacteria. Antigens are *B. cellulosilyticus* (positive control), RD014212 (*Catabacter sp.*), RD014215 (*Clostridium sp.*), RD014238 (*Flavonifractor sp.*), RD014241 (*Intestinimonas sp.*), RD014243 (*Peptoniphilus sp.*), RD014249 (*Faecalicoccus sp.*), RD014895 (*Mediterraneibacter sp.*), *S. cerevisiae*, HEK293T cells, and BSA. Bars indicate the average of triplicate measurements ± standard deviations.

### 3.6. Kinetics analysis of Fab

The binding properties of the purified Fab expressed in the *Brevibacillus choshinensis* expression system were examined by the biolayer interferometry method using inactivated *B. cellulosilyticus* cells as the antigens, which were immobilized on APS sensors, followed by association with Fab antibodies at various concentrations. After dissociation, the sensorgrams were analyzed, resulting in *K*_D_ values of 2.41 nM, as shown in Fig. 7. These data indicated that the mAb developed by Ecobody technology and produced in the *Brevibacillus choshinensis* expression system is fully functional with a high affinity.

**Fig. 7.**
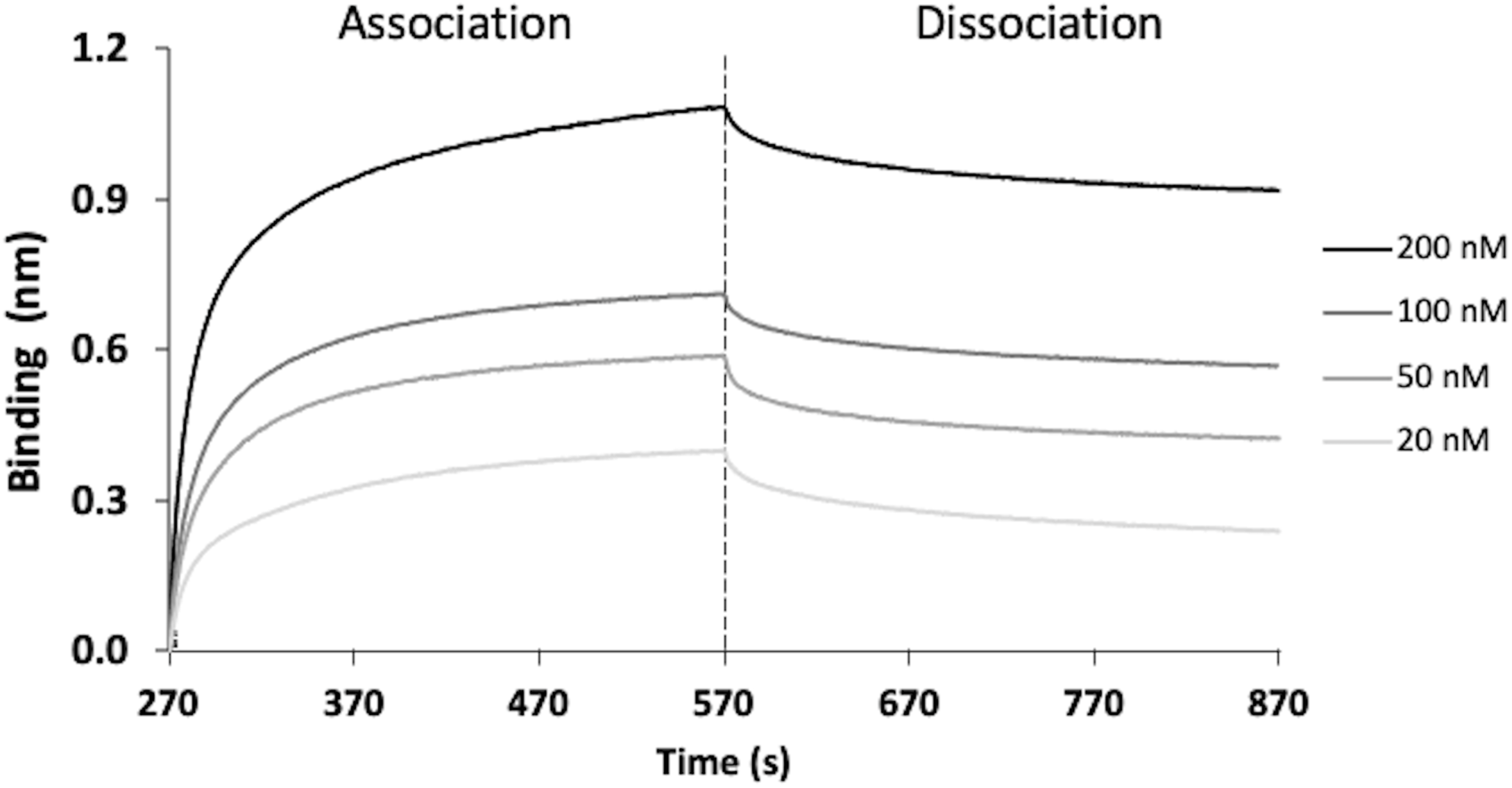
BLItz sensorgram at different concentrations of the Fab produced in the *Brevibacillus choshinensis* expression system against inactivated cells of *B. cellulosilyticus* (OD_600_ = 1.0)

## 4. Discussion

In this study, we successfully obtained and investigated intestinal bacteria-binding monoclonal antibodies derived from single B cells of a non-immunized rabbit using Ecobody technology coupled with the *Brevibacillus choshinensis* expression system. The sequencing of 16S rDNA was performed to validate the presence of *B. cellulosilyticus* as the targeted bacterium within the rabbit intestinal microbiota. We investigated several phyla from the rabbit cecum, including Firmicutes, Bacteroidetes, Verrucomicrobia, Actinobacteria, and Euryarchaeota. Firmicutes was the most dominant phylum detected in the rabbit cecum in agreement with previous results from Velasco-Galilea [1], followed by Bacteroidetes, which includes the genus *Bacteroides*. Other phyla, such as Verrucomicrobia, Actinobacteria, and Euryarchaeota were found in relatively small abundance but still contributed to shaping the diversity and stability of the microbial community in the rabbit cecum. Although the number of phyla detected in this study was fewer than in the results of a previous study [28], our data confirmed that *Bacteroides* is a part of the rabbit intestinal microbiota.

Following the validation of the presence of *B. cellulosilyticus* within the rabbit intestinal microbiota, we utilized Ecobody technology for rapid screening of mAbs targeting intestinal bacteria from single B cells, from which antigen-specific B cells were obtained using the inactivated *B. cellulosilyticus* cells as the antigen. The Lc and Hc genes were separately amplified from single B cells and cloned into the expression vectors harbouring the SKIK tag and leucine zipper (LZ) tags for cell-free protein synthesis (CFPS), as shown in Fig. 2. Moreover, the number of clones from the same cells was very limited and only IgM clones could be obtained, indicating that single B cells originating from the peripheral blood of a non-immunized rabbit have low diversification. Consequently, we combined the Lc and Hc originating from different cells in Fab expression. The ELISA result indicated that most of the synthesized Fab antibodies had binding activity toward *B. cellulosilyticus* and did not cross-react with BSA (Fig. 4). This is probably because non-specific B cells that bound to BSA were successfully removed during the selection process and solely remaining antigen-specific B cells that bound to *B. cellulosilyticus*. In addition, the utilization of this simply subtractive screening technique proves to be a highly efficient method for isolating antigen-specific B cells [17]. The success of antigen-specific antibody acquisition is consistent with previous studies [16, 18, 29, 30].

One mAb clone (L86M86) consisting of the cognate pairing of Lc and Hc was selected and further produced in the *Brevibacillus choshinensis* expression system. Most of the target proteins were secreted into the medium culture and easily harvested for purification steps. We selected proline, Arg-HCl, and MgSO_4_ as supplementary compounds because their function as folding auxiliary molecules is consistent with previous reports [23–26]. We observed that the addition of these molecules also enhanced bacterial growth and Fab expression in *Brevibacillus choshinensis* as previously reported in the reference [23]. However, the effect of supplementation of these molecules appears to be different in some cases of recombinant protein expression. Therefore, it is necessary to optimize the composition of the culture medium before large-scale production.

In the present study, we have successfully expressed mAb clone 86 in the active form of Fab and yielded about ∼2.0 mg of the purified Fab antibody from *Brevibacillus choshinensis* per Liter of the culture in the supplemented TM medium using 96-deep-well plate culture (Supplementary Table 3). The purified Fab that was obtained in this study was still lower than the results from a previous study [23], although the direct comparison is not feasible because of the different genes used. To date, 100 mg/L culture of anti-urokinase-type plasminogen activator Fab in *Bacillus brevis* [31] and 145 mg/L of Trastuzumab Fab (without His-tag) in *Brevibacillus choshinensis* [32] were reported. Gram-positive bacteria effectively secrete several proteins into the culture as they lack an outer membrane, which acts as a barrier in Gram-negative bacteria, preventing the retention of secretory proteins in the periplasmic [33].

The purified Fab was then tested against several Gram-negative and Gram-positive bacteria for investigation of cross-reactivity. Through rigorous testing, the ELISA signal revealed that Fab could bind to both Gram-negative [Fig. 6a] and Gram-positive bacteria [Fig. 6b] but did not bind toward *S. cerevisiae* and HEK293T cells, indicating the specificity for intestinal bacteria-associated antigens. Interestingly, our data showed that the LPS derived from Gram-negative bacteria is one of the antigens targeted by the obtained mAb due to its binding activity against the LPS [Fig. 6a], only slightly lower than the binding activity toward the inactivated bacterial cells. The binding activity against Gram-positive bacteria is likely due to other potential antigens recognized by antibodies, including peptidoglycan (PG), which is generally found on the surface of Gram-positive bacteria. Nevertheless, further analysis using extracted peptidoglycans as the antigen is required to prove our hypothesis. Previous studies reported cross-species reactivity of the mAbs recognizing conserved antigens that are structurally identical antigens shared among different members of intestinal bacteria, such as lipopolysaccharide (LPS) and peptidoglycan (PG). Bacterial glycans inducing the mucosal immune system are typically T-independent antigens that tend to produce mainly low-affinity and polyreactive IgM antibodies [13, 34, 35].

The exposure of multiple bacterial antigens promotes the formation of cross-reactive antibodies in rabbit intestines. The cross-species reactivity selection is likely a result of a gradual accumulation of somatic mutations through continuous affinity maturation in germinal centres (GCs), especially in Peyer’s patches of the GALT [11]. This process facilitates the development of broad yet specific antibody responses, allowing for rapid adaptation to significant changes in microbiota composition and serving as a protective agent against broadly bacterial infection as previously reported [36–39]. We therefore suggest that the cross-species reactivity of antibodies is a common feature of antibody-microbiota interactions in the intestine.

Using Ecobody technology, we succeeded in obtaining and investigating monoclonal antibodies targeting intestinal bacteria without rabbit immunization coupled with the *Brevibacillus choshinensis* expression system for large-scale antibody production, demonstrating that the purified Fab possesses a high affinity (*K*_D_ = 2.41 nM) to the target bacterium (Fig. 7) and a broad reactivity toward several intestinal bacteria. Ecobody technology significantly reduces the time needed for antibody screening and production compared to traditional hybridoma technology. Our data prove the applicability of Ecobody technology in obtaining and generating mAbs targeting conserved antigens from intestinal bacteria that might not elicit a strong immune response in conventional immunization protocols. The capability of this method to obtain and produce different types of mAb clones needs to be further investigated, but we expect that Ecobody technology coupled with the *Brevibacillus choshinensis* expression system offers an alternative technique for rapid screening and large-scale production of monoclonal antibodies at low cost.

## 5. Conclusion

In conclusion, intestinal bacteria-binding mAbs derived from rabbit single B cells were successfully obtained and investigated by Ecobody technology coupled with the *Brevibacillus choshinensis* expression system, demonstrating a high affinity. The Fab revealed the cross-reactivity toward several intestinal bacteria but did not bind to unrelated antigens, suggesting the specificity toward intestinal bacteria-associated antigens. We consider Ecobody technology coupled with the *Brevibacillus choshinensis* expression system to be convenient and applicable for a high-throughput method of mAb screening and production that may contribute to antibody development for various purposes in research and industry.

## Acknowledgments

This research was financially supported in part by Grants-in-Aid for Scientific Research (No. 22520782 19105921) of the Japan Society for the Promotion of Science (JSPS), and by the Institute for Fermentation, Osaka IFO): IFO research grant G-2022-3-021. Khairil Anwar is supported by the scholarship from The Ministry of Education, Culture, Sports, Science and Technology (MEXT) from Japanese Government and Graduate Program of Transformative Chem-Bio Research (GTR), Nagoya University. Daffa Sean Adinegoro gratefully acknowledges to GTR Program. Daffa Sean Adinegoro and Monami Kihara are supported by THERS Make New Standards Program for Next Generation Researchers. The authors thank Gene Frontier Corporation, Japan, for providing PURE*frex* components. We are grateful to Prof. Toru Suzuki and all the staff at Gifu University’s Next Generation Sequencer Service for 16S rDNA analysis.

