## Supplementary Data for "Investigation of Intestinal Bacteria-binding Monoclonal Antibodies Derived from Rabbit Single B Cells"

**Supplementary Table 1. Specific DNA primers for IgA antibody**

| Description | Target gene | Sequence (5' to 3') |
|---|---|---|
| Reverse transcription | Hc (IgA) | GAAGTTATAGACTCTCACGTTCTC |
| | | GGGAAGATCACGTCCCCCTC |
| | | TAGGATCAGCTCGCTGCATG |
| | | TAGGCTCAGCAGGCTGCATG |
| | | GGGAAGGTCACGTTCTCCCC |
| | | GGGAAGATCACGCCCCCCTC |
| First PCR | Hc (IgA) | GTTCTCCCCACTGACATCCCAGCTCAC |
| | | GCTGCCGTTCCAGCTCACATTC |
| | | TACGGGCCTGAAGTGCCAG |
| | | TTCAGGCCTGAAGTGCCAG |
| | | GCTGCCGTTCCATGTCACATTC |
| | | GCTGATGGTCCAGTTCACATTC |
| Second PCR | Hc (IgA) | TCCGCTTACGTCCCAACTTACACTCAGAGGGTCCAGTGGGAAG |
| | | TCCGCTTACGTCCCAACTTACACGCAGAGGGCCCAGGGGGAAG |
| | | TCCGCTTACGTCCCAACTTACACGCAGAGGGCTCGGGGGGAAG |
| | | TCCGCTTACGTCCCAACTTACACTCAGAGGGCCCGGGGGGAAG |
| | | TCCGCTTACGTCCCAACTTACACTCAGAGGGCCCAGGGGGAAG |
| | | ACTACCATTCCAACTGACGTTCAGAGGGCCCAGGGGGAAGAA |

**Supplementary Table 2. Primers for construction of pNCMO2-Fab clone 86 (L86M86) for the *Brevibacillus choshinensis* expression expression**

| Target | Name | Sequence (5’ to 3’) |
|---|---|---|
| Signal 3 | pBIC-Signal3-F | ATGTCAATTTCGGTAAGATTCAAGAG |
| | pBIC-Signal3-R | AGCGAATGCGGAACTTGGTAC |
| Signal 4 | pBIC-Signal4-F | ATGAAAAAAAGAAGGGTCGTTAACAG |
| | pBIC-Signal4-R | AGCGAAAGCCATGGGAGC |
| pNCMO2-vector | pNCMO2-Sig3-inv-R | TCTTACCGAAATTGACATAGCGCGTGTCACCTCCTTTC |
| | pNCMO2-Sig3-inv-F | CACCATCACCATCACCATTAAAGAGGAGGAGAACACAAGGTC |
| | pNCMO2-Sig4-inv-R | GACCCTTCTTTTTTTCATGACCTTGTGTTCTCCTCCTCTTTAATG |
| Lc86 | Sig3-LcK86-F | CCAAGTTCCGCATTCGCTGACATTGTGATGACCCAGACTCC |
| | Sig3-LcK86-R | TTAATGGTGATGGTGATGGTGGCTCCCACCACCGCCAC |
| Hc86 | Sig4-HcIgM86-F | GCTCCCATGGCTTTCGCTCAGTCGGTGAAGGAGTCCG |
| | Sig4-HcIgM86-R | TTAGTGATGGTGATGGTGATGCGGAAAGCTAACACGCAGATC |

**Supplementary Table 3. Recovery ratio of *Brevibacillus Choshinensis* expressed proteins**

| Purification steps | Fab amount (mg) | Yield (%) |
|---|---|---|
| Culture supernatant (222 mL) | 2.71 | 100.0 |
| Ni-column | 1.26 | 46.5 |
| SEC column | 0.57 | 21.0 |

The Fab antibody was expressed in the *Brevibacillus choshinensis* expression system using 288 mL of TM media supplemented with 200 mM arginine hydrochloride, 10 g/L of proline, 60 mM $MgSO_4$, and 50 μg/mL neomycin. The culture was performed using 96-deep well plates at 30 ºC for 55 hours with reciprocal shaking (1000 rpm). Proteins were harvested from the culture supernatant and regarded as 100% in recovery ratio.

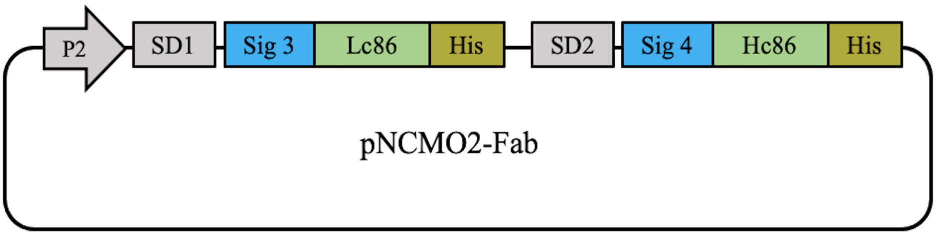


**Supplementary Fig. S1. Plasmid designed for Fab expression in the *Brevibacillus choshinensis* expression system**.

The pNCMO2-Fab plasmid was constructed to harbour tandem Lc and Hc expression cassettes on a single plasmid. The target genes was inserted after Shine-Dalgarno sites (SD), the light chain (Lc) after SD1 and heavy chain (Hc) after SD2. P2 promoter is the main promoter derived from the cell wall protein synthesis of *Brevibacillus*. Secretion signals (Signal 3 and Signal 4) are fused at the N-terminus of the Lc and Hc, respectively. A polyhistidine tag (6×his tag) was inserted at the C-terminus of both Lc and Hc, respectively.

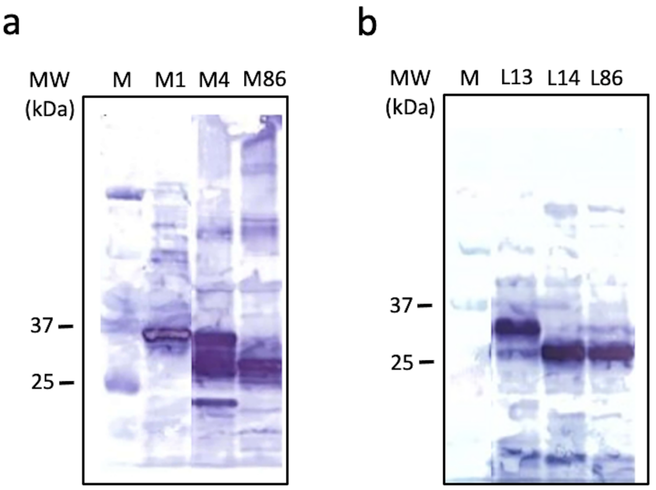


**Supplementary Fig. S2. Western blotting analysis of Fab-LZ with N-terminal SKIK tag synthesized by CFPS**

a) Western blotting analysis of the Hc detected using anti-HA tag-HRP conjugated antibodies. M: Prestained protein standards (Bio-Rad), M1: Hc IgM clone No. 1, M4: Hc IgM clone No. 4, and M86: Hc IgM clone No. 86.

b) Western blotting analysis of the Lc detected using Anti-PA tag-HRP conjugated antibodies. M: Prestained protein standards (Bio-Rad), L13: Lc clone No. 13, L14: Lc clone No. 14, and L86: Lc clone No. 86.
